# Biological Insights Knowledge Graph: an integrated knowledge graph to support drug development

**DOI:** 10.1101/2021.10.28.466262

**Authors:** David Geleta, Andriy Nikolov, Gavin Edwards, Anna Gogleva, Richard Jackson, Erik Jansson, Andrej Lamov, Sebastian Nilsson, Marina Pettersson, Vladimir Poroshin, Benedek Rozemberczki, Timothy Scrivener, Michael Ughetto, Eliseo Papa

**Affiliations:** Research D&A, R&D IT, AstraZeneca

## Abstract

The use of knowledge graphs as a data source for machine learning methods to solve complex problems in life sciences has rapidly become popular in recent years. Our Biological Insights Knowledge Graph (BIKG) combines relevant data for drug development from public as well as internal data sources to provide insights for a range of tasks: from identifying new targets to repurposing existing drugs. Besides the common requirements to organisational knowledge graphs such as being able to capture the domain precisely and give the users the ability to search and query the data, the focus on handling multiple use cases and supporting use case-specific machine learning models presents additional challenges: the data models must also be streamlined for the performance of downstream tasks; graph content must be easily customisable for different use cases; different projections of the graph content are required to support a wider range of different consumption modes. In this paper we describe our main design choices in implementation of the BIKG graph and discuss different aspects of its life cycle: from graph construction to exploitation.

## 1 Introduction

Recent years have seen rapid growth in the popularity of knowledge graphs in the life sciences domain. Knowledge graphs often serve as a backbone for data integration within organisations, providing a common representation structure which enables querying across data sources. With the recent advances in machine learning (ML), knowledge graphs gained one more important purpose: to serve as training data for ML models, and graph machine learning models in particular. Their newfound use as training data has to be taken into account when constructing knowledge graphs and influences core design choices. For example, while a very expressive schema can be able to capture the most fine-grained aspects of domain data, it can at the same time hinder the application of machine learning due to scalability problems and diffusion of signal. Similarly, use as ML training data necessitates support for different modalities of data usage, beyond structured queries.

In this paper we describe the Biological Insights Knowledge Graph (BIKG): an AstraZeneca project aimed at building a knowledge graph combining both public and internal data to facilitate knowledge discovery using machine learning. BIKG integrates knowledge from heterogeneous data sources including public databases like ChEMBL [14] or Ensembl [50], information extracted from full-text publications using Natural Language Processing (NLP) techniques, as well as diverse proprietary datasets collected as part of AstraZeneca drug development process and biological experimentation.

The rest of the paper is organised as follows:

- Section 2 discusses the main requirements to the BIKG knowledge graph and compares it with existing similar initiatives.
- Section 3 describes the process of building the knowledge graph by integrating heterogeneous sources.
- Section 4 describes different modalities of user interaction with BIKG data.
- Section 5 outlines the main applications of machine learning algorithms to graph data.
- Section 6 introduces the initial practical use case scenarios exploiting graph data.
- Section 7 discusses the main lessons learnt and lists directions for the future work.

## 2 Motivation and Related Work

The life sciences domain has been an early adopter of ontologies and linked data technologies. The primary motivation, given the vast amount of knowledge to capture, was the need for standardisation of the vocabularies and taxonomies. This was essential for integration of data within and across large organisations and enabling data access. For this reason, the design of graph datasets focused on high granularity of the models and ability to capture the complexity of the domain in the most precise way. Such ontologies included, for example, the Gene Ontology (GO) [1] for gene functions, the Human Disease Ontology (DO) [22] for capturing different disease classifications, or BioPax [11] for pathway-related information.

With the growing number of datasets capturing separate domains, the focus shifted towards achieving interoperability and building integrated datasets that could serve as reference data sources across multiple interconnected topics. These included both “vertical” ontology integration of semantic alignment of multiple ontologies via common foundational schemas (e.g., the OBO Foundry [42]) and “horizontal” integration focusing on the data-level fusion of separate datasets into large interconnected graphs covering many domains that could be transparently queried (e.g., the Bio2RDF [3] triple database). Standardizing data structure and naming conventions helped to improve reusability of scientific data according to the FAIR data principles [47], making knowledge graphs a backbone for large-scale integration of data.

More recently, with the advances in network medicine approaches [2] and in the area of AI in general, another role for knowledge graphs is quickly gaining importance, namely, facilitation of machine learning. Machine learning methods on graphs often help to overcome sparsity of reliable data and reduce the need for expensive and time-consuming experiments by pre-selecting the most promising candidates for manual verification. Graph machine learning has been applied to a number of tasks [13] such as target identification [35], drug repurposing [16], or predicting polypharmacy side effects [53].

With the development of machine learning algorithms processing life science knowledge graphs, more work has focused on constructing integrated knowledge graphs adapted to support such algorithms. There are several recent initiatives focusing on building knowledge graphs combining information from multiple data sources to use them for training machine learning models. For example, OpenBIOLink [5] was developed to provide a common benchmark to evaluate machine learning algorithms on life-science-related tasks. Drug Repurposing Knowledge Graph (DRKG) [19, 48] specifically combines the most relevant sources to support the drug repurposing task. CKG [38] focuses on bringing together multi-omics data to support precision medicine.

Constructing knowledge graphs to support graph analytics and machine learning has its own set of requirements that do not completely match the scenarios where the primary use of the graph is to enable complex cross-source querying. For example, a very precise and detailed data model can make it hard for an algorithm to learn essential relations between nodes, if they are not connected directly, but separated by several hops. High model granularity can also increase the size of the graph and make some complex models expensive to train.

The usability expectations also differ in this case. While the abilities to search and formulate expressive queries are still important, the possibility to customise and manipulate the graph for different use cases becomes particularly valuable. Depending on the task (e.g., drug repurposing or target selection) or the domain (e.g., specific disease like asthma), the users should be able to extract a custom relevant subgraph to apply statistical methods and train models. In general, it is important to combine for analysis both public reference datasets capturing state-of-the-art knowledge (e.g., Ensembl [50] for genes or ChEMBL [14] for drug compounds) as well as internal proprietary data generated inside the organisation over many years. Moreover, researchers often maintain their own task-specific datasets which they want to include in analysis, together with large-scale reference data sources. For this reason, integrating new data into the graph should be made as easy as possible, as well as the filtering of the graph according to custom criteria. Finally, the graph has to be easily consumable as input by popular machine learning software libraries.

With these requirements in mind, we developed the BIKG (Biological Insights Knowledge Graph): an internal AstraZeneca knowledge graph aimed at supporting analytics and machine learning tasks to help drug development. The graph construction process focuses on unifying the structure of multiple source datasets and the flexibility of the workflow: enabling quick incorporation of feedback and “democratisation” of the data integration by allowing the end users to bring in their own tasks-specific data. Multiple graph usage modalities, in turn, provide alternative ways of interacting with the graph optimised for different end-user tasks.

The BIKG development process involves supporting two major stages:

- *Graph construction*, which involves bringing together diverse data sources and performing common data integration tasks to produce the graph and provide multiple access options.
- *Graph utilization*, which involves supporting the end users with applying machine learning techniques to solve use case tasks.

## 3 Knowledge Graph Construction

The graph build pipeline integrates internal and external data, implemented in the cloud as a secure and scalable pipeline architecture, to create a consistent and coherent knowledge graph that can be used for applying machine learning algorithms and gaining new insights.

This section is organised as follows: Section 3.1 details the data sources used in the graph, and outlines how data can be added to future graph builds or used with an existing graph version. Section 3.2 introduces the low level schema (unified data model). Section 3.4 talks about how graph compression is used to managed the size. Section 3.3 explains how we make the graph accessible for a wide range of audience by producing several formats, i.e. graph *projections*. Section 3.5 introduces the high level schema, i.e. the upper level ontology. Section 3.6 describes node identity resolution. Section 3.7 shows how the graph quality is evaluated. The main pipeline steps are introduced in Section 3.8. Finally, Section 3.9 mentions the technology stack used in the pipeline.

### 3.1 Data sources

The BIKG graph integrates information from multiple sources. The latest graph combines over 50 years of biomedical information into a single resource, consisting of 10.9m nodes (of 22 types) and over 118 million unique edges (of 59 types, forming 398 different triples) as shown in Table 1. Some of the ingested data sets are themselves integrate a number of data sources, such as *Hetionet* [18] *and Opentargets* [27].

**Table 1.** The graph content summary table. provides a flavour for its size and compositional variety.

| Summary | Count |
| --- | --- |
| Nodes | 11m |
| Node types | 22 |
| Node contexts keys | 276 |
| Edges (collapsed) | 118m |
| Edges (uncollapsed) | 1189m |
| Edge types | 59 |
| Edge evidence keys | 154 |
| Triple types | 398 |
| Ingested data sets | 39+ |

At the core of the graph are reference datasets describing different topics of the drug discovery domain, such as genes, compounds, and diseases. These reference datasets are enriched with relations automatically extracted from literature using state-of-the-art natural language processing. Finally, in order to adjust to the needs of the end users and their tasks, the users have the ability to enrich the graph with their task-specific data. Using BIKG allows users to connect to *internal company data* and gain a unique advantage with respect to openly available graph datasets.

#### 3.1.1 Backbone: static reference datasets

Standard datasets describing specific domains constitute the “backbone” of the BIKG graph: they both define the high-level structure for representation of each respective domain and the identification conventions for domain entities. This includes popular standard datasets commonly used in drug discovery [4], such as Ensembl [50] for genes, ChEMBL [14] for drug compounds, Mondo [32] for diseases, the Gene Ontology [1], and others. To model each domain of interest in the most complete way, in cases where there exist several alternative reference sources, they were integrated together and merged using mappings between the corresponding identifiers: e.g., having ChEMBL identifiers as canonical ones for drug compounds and expanding them with complementary external (such as PubChem [23]) as well as internal ones (e.g., covering experimental formulas not yet indexed by public repositories). Apart from the actual content data, integration of mapping sets provided by different sources represents added value in itself, allowing end users to switch quickly between the naming schema of their use case-specific source and the canonical one used by the graph.

#### 3.1.2 NLP for graph population

While the data imported from various structured sources provides the most reliable part of the graph, it is enriched using large-scale relation extraction from free-text sources such as scientific literature from PubMed^1^. The NLP aspects of BIKG is based on a series of pipelines, ranging from simple entity co-occurrence and traditional rule based dependency parsing, to state-of-the-art relationship classification with RBERT [49] and open information extraction with OpenIE6 [26] neural information extraction system. In terms of quantity, this NLP-extracted data constitutes the largest component of the graph, providing around 80% of graph edges. Despite inherent uncertainty associated with these edges due to potential NLP errors, when aggregated, they provide clear added value for the output quality of machine learning models: e.g., NLP-extracted edges were found to be the most informative in a link prediction benchmark task focusing on protein-protein interaction (PPI) network population.

Extracting a large amount of structured information from biomedical literature remains a very complex task. The language used to describe biological entities and relationships varies across temporal and geographic dimensions, and is subject to various phenomena in language evolution (for instance, neologisms, synonyms, multi-entity constructs etc).

Progress in developing a generalised knowledge base population solution is hampered by:

- Lack of sufficiently diverse training data covering the breath and depth of biomedical knowledge
- Biases in existing training data sets (such as only covering a subset of diseases)
- Uncertainty/fluidity in our understanding of disease mechanisms, resulting in inconsistent language usage over time
- Contradictory evidence and or irreproducible results
- Uncertainly in the interpretation of data, often resulting in authors hedging the assertions they make
- Biases in literature to only report positive findings
- Poor generalisability of ML models, research findings are generally overfit to small datasets and not suitable for production offerings

Our NLP data is derived from a multi stage pipeline:

1. Abbreviation Expansion - In order to improve our named entity recognition, we preprocess text data to expand abbreviations. [39]
2. Named Entity Recognition - to identify graph entities, we use the commercial Termite Tagger from Scibite. We supplement this with mutation information, based upon a modified form of SETH.
3. NER Post Processing - in this stage, we normalise entity information from different NER tools back to a standard data class; enrich entities with BIKG ids where possible; and perform entity linking on concepts that otherwise lack this information
4. Relationship Extraction

Our relationship extraction engine is based on three techniques:

- a rules based dependency parsing application, based on the open source LINK software, and
- relationship classification using a neural network based upon BioBERT [29] and the RBERT [49] classification head.
- Open Information Extraction using the OIE6 software [26].

We limit the overall complexity of the input dataset by excluding those sentences containing more than 30 entities from any downstream processing. Complex sentences with more than 12 entities are not fed to our neural network inference engine.

##### LINK

Modern dependency parsers are incredibly accurate. However, while they are informative about the syntactic structure of sentences, they offer no guidance on how such structure should be manipulated in order to extract triples. Our current work in this area involves the refactoring of the LINK NLP pipeline [34] to make use of our commercial NER tool, Termite [40], and the subsequent crafting of rules to produce triples suitable for ingestion into the graph.

An advantage of dependency parsing approaches is that they are able to predict the type of relation (verb) between two entities, meaning that the results can be mapped to an ontology. The drawback is that they tend to be more error prone than neural network based methods.

##### RBERT

Taking inspiration from the latest wave of attention based transformer NN, we have written our own SOTA relationship classifier implementation based upon the work of Wu and He. [49] The pipelines achieved a new state-of-art (SOTA)) benchmark on the popular SemEval 2010 Task 8 [36] relationship classification.

Our RBERT model is more powerful than LINK, but suffers from the significant disadvantage of only being able to suggest associations between two entities, as opposed to detecting the verb describing the relationship.

Our deployed model is trained to predict binary relationships between the following entity types:

- Gene Target—Gene Target
- Gene Target—Compound

To give an indication of performance, we use the BioInfer [37] and BioRelex [21] datasets. A thorough analysis of the performance of this pipeline is included elsewhere [20].

#### 3.1.3 User-contributed data

One of the main requirements of BIKG is the possibility to integrate internal use case-specific datasets on demand. Given the multitude of use cases, a purely centralised approach to data addition has limitations, because constant need to process new datasets creates a bottleneck on the side of the graph engineering team, while it is the end users and data owners who have the best knowledge of the data and the use case needs. For this reason, the approach chosen in BIKG is to enable and support the end users in integrating the data they need. This task was made easier by the fact that the majority of graph users are bioinformaticians who are well familiar with script coding as well as tabular data manipulation techniques.

The *Bring Your Own Data (BYOD)* module, which is part of the Python API (used for accessing the graph, see Section 5.4), aids the ends users in extending the graph. This provides a command-line interface, code templates as well as tutorial notebooks, to aid users to transform their data into a pre-defined tabular format suitable for automated insertion into the graph. As the data might be syntactically well formed and its semantics need to conform to the BIKG Upper Level Ontology (Section 3.5), the users are also provided with a data verification tool, which checks the data format, relational structure, as well as the naming conventions for node identifiers. Using this API users can parse and integrate their own data to a copy of the graph, and can also share datasets to be integrated into to central build.

### 3.2 Unified data model

Similarly to the *Microsoft Academic Graph (MAG)* [41], *the data is modelled and distributed as a set of tables. In the BIKG pipeline the graph data is processed and stored in parquet* files^2^. There are two main tables, nodes (including mappings) and edges, described in Section 3.2.1 and Section 3.2.2, respectively. Both tables contain *standard columns* (representing mandatory table fields) and *dynamic columns* for representing the different node or edge background information found in the ingested sources.

#### 3.2.1 Nodes

A node is defined as the tuple *node* ⟨ *id, label, type, context*(*c*_*i*_…*c*_*j*_)⟩, where the fields are defined as follows:

- *id*: the preferred identifier for a given node, adhering to a unified ID schema (*DATA_SET_ID* : *CODE*, which is the internal node ID where the original ID is prefixed with a namespace to capture provenance e.g. *ENSEMBL* : *ENSG*0001 for the Ensembl node *ENSG*0001);
- *label*: the preferred label for the node, for example for genes often the gene symbol label is used;
- *type*: the classification of the node, this must conform to the ULO (see Section 3.5).

In parsed sources node tables also have a *provenance* column to aid the build pipeline identity resolution step (see Section 3.6) in finding a canonical node for duplicate node instances (i.e. nodes with the with the same *id* appearing in different parsed sources). In addition to the above described *standard columns* that must be present in all parsed sources, the node table may have a set of context columns, *context*(*c*_*i*_…*c*_*j*_). These columns record background information about the node such as labels, types or other data that may be a potential feature in ML training:

- *context*.*labels*: All labels (preferred, synonym, description, definition, notes etc) associated with the concept
- *context*.*types*: All types that are associated with a given node ID over all of its instances in the loaded data; e.g. X can be a Disease but also a Side Effect, storing these enables for better filtering. The *type* value is a member of this set.

Node mappings are stored as a separate table in the build process, but materialised in graph projections for more convenient usage.

#### 3.2.2 Edges

The tuple *edge* ⟨*source_id, target_id, relation, provenance, evidence*(*e*_*i*_…*e*_*j*_)⟩ defines an edge where:

- *edge_id*: the edge identifier is the base64 encoding of the fields *source_id*,
- *target_id, relation*, and *provenance* (note that this is not necessarily unique in a data set prior to graph compression (see Section 3.4);
- *source_id, target_id*: identifiers of the nodes connected by an edge;
- *relation*: the type of the relationship between the source and the target node;
- *provenance*: the source data set identifier;
- *publications*: most BIKG edges have referencing publication or patent as underpinning evidence, these are aggregated from typically several columns;
- *evidence*(*e*_*i*_…*e*_*j*_): zero or more meta data columns associated with the edge.

In a compressed edge table (see Section 3.4), standard edge columns (*source_id, target_id, relation, provenance*) are the same, but the evidences are merged.

### 3.3 Graph projections

The graph is built using several columnar data tables (node, mapping and edge table) that enable fast processing as well as reduce overall data size. The resulting output contains all data, such as mappings and alternative labels for nodes, as well as a large number of features (contextual attributes for nodes, and underpinning evidence for edges). In this form, the graph is not user friendly because it contains billions of rows and hundreds of columns. Unless users are equipped with big data processing infrastructure, even simple tasks (for example querying the graph or creating edge sets) can pose a challenge. In order to support users and to make the graph more accessible for a wide range of audience within the company, several formats, denoted as *graph projections*, are produced by the pipeline. Section 4 describes how these projections are used. Projections may contain materialised data (e.g. node mappings are inferred and materialised in the nodes table), may differ in data structure (in the BIKG projection, the edge table is split into two tables), or uses a dedicate format (e.g. RDF for triple stores). The following projections are produced:

- The *BIKG* projection is the main distribution artefact, i.e. the source of truth, which is used by the Python API (see Section 5.4). This is a set of parquet files, where the edge table into two tables, one for the standard columns representing the edge (*source_id, relation, target_id, provenance*) and another containing all feature columns (*evidence*(*e*_*i*_…*e*_*j*_)); as a result users can pick and choose only desired features, ultimately reducing data size. The projection uses compressed edges (see Section 3.4).
- The *BROWSER* (Section 4.1) projection is also serialised as parquet files, but it contains the uncompressed edges in order to aid indexing and data presentation, where standard edge information is merged with the feature columns.
- The *RDF* 4.2 projection (N-Triple files^3^) and the *Neo4j* 4.2 projections (CSV files) both use the compressed edges. In the RDF format, nodes are materialised as classes, edges as triples, where the corresponding edge features are represented as triple annotations. In addition to the aforementioned graph database formats, another RDF projection is generated; this presents the high level schema, graph meta data and analysis (such as node and triple type counts) to support graphical graph exploration [30].

### 3.4 Graph compression

The graph is compressed and filtered to limit its size. This is necessary because the raw data set is quite large (several terabytes); the data is noisy; also the purpose of the graph is not information retrieval but facilitating signal propagation, hence not all ingested data is preserved in the main graph projection (however, note that relevant data, e.g. publication references, is stored prior to the graph build and is made available separately as needed by users).

The graph is compressed by *merging duplicate edges*. Two edges are considered duplicates if they shares the same *source_id, target_id, relation, provenance* attributes, i.e. they have the same source and target node identifier reference, relation type and edge provenance, hence the only distinguishing feature is the difference in evidence attributes (e.g. publication source, experimental evidence score value etc.). When merging edges, the evidences are retained. However, this set can be rather large due to: skewed literature data (for example, several trivial edges are parsed from many publications, resulting in millions of noisy triples); and overlapping data source content. Therefore, to limit the size of the graph, the number of merged edges for each *edge duplication case are limited*. This reduces noise and retains most relevant information. The number of duplicate edges is retained with each merged edge.

### 3.5 Upper Level Ontology (ULO)

Different source datasets of BIKG use their own vocabularies to represent the same domains and model different aspects of the data with different levels of granularity. For this reason, a unification of models is required to produce an integrated graph with a common structure. The Upper-Level Ontology (ULO) of BIKG (see Figure 2) serves this purpose by defining a common schema over integrated data sources. The second goal of the ULO ontology is to define common data constraints to verify the consistency of integrated data.

**Figure 1.**
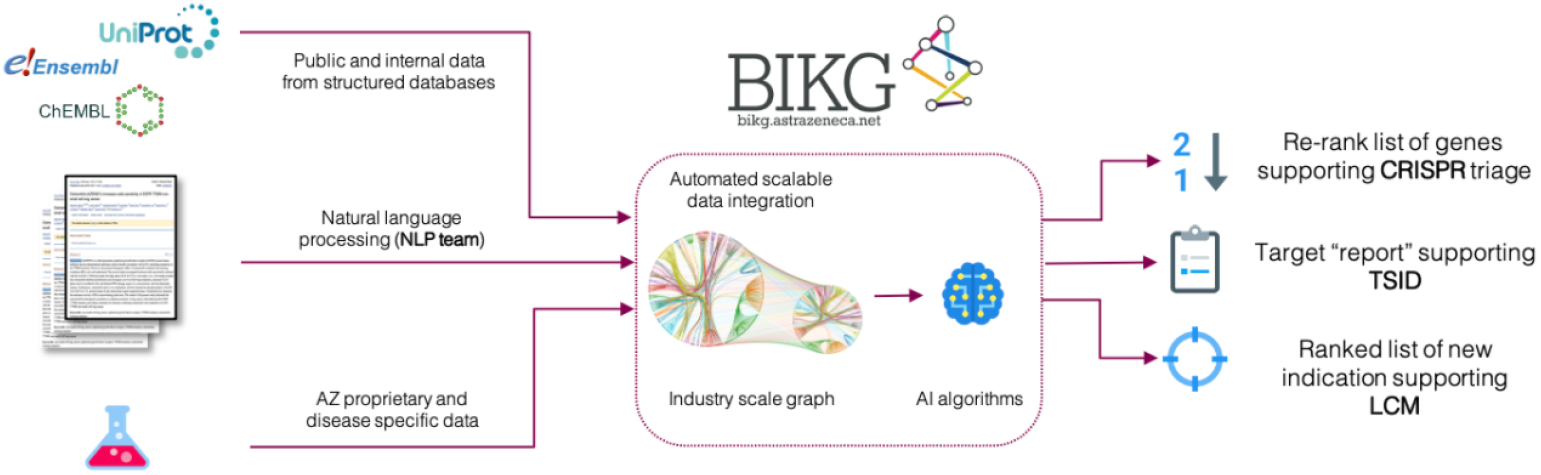
The Biological Insights Knowledge Graph project overview: data types, graph build, and example use cases.

**Figure 2.**
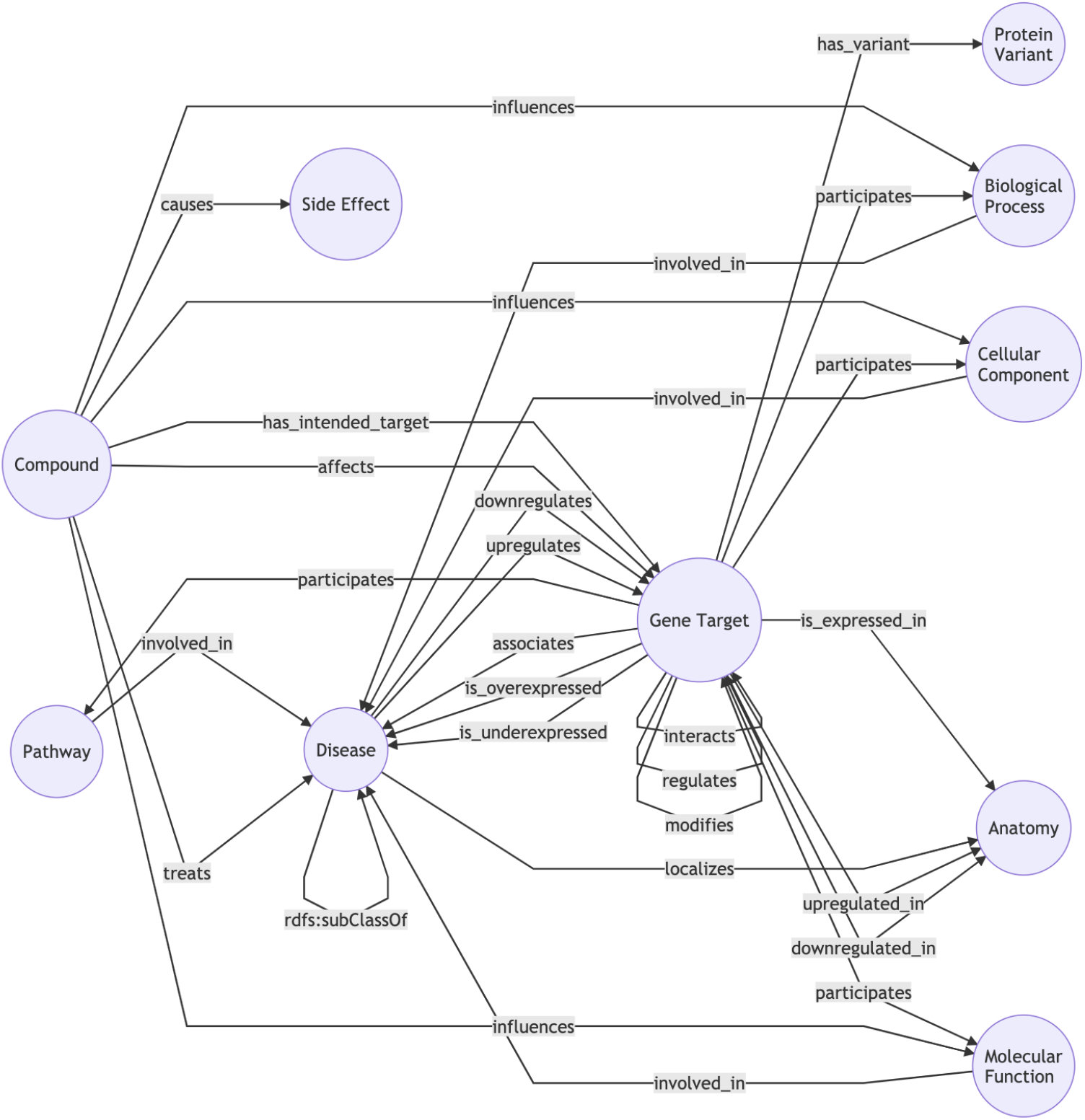
Core fragment of the BIKG Upper Level Ontology

ULO aims at representing the node and edge types at the level of granularity that enables best signal propagation while, at the same time, providing sufficient coverage for representing the domain area. At the core of the ULO ontology are the most relevant concepts of the drug development domain: Compound, Gene Target, and Disease and relations between them. Design choices were motivated by the needs to achieve uniform representation of the source data and to facilitate the training of machine learning algorithms. For example, the notion of a *target* is a key concept in drug development, usually representing a protein affected by a drug compound and encoded by some gene. However, different sources use different conventions for modelling targets: some refer to targets using the actual protein identifiers (e.g., codes from the UniProt database [46]), while many others use the identifier of the encoding gene (such as an Ensembl code [50]). Given that the gene-protein relations are in itself indirect (representing the gene *→* transcript *→* protein chain) and not always 1:1, integration of different sources while preserving their full level of detail would necessarily break the uniformity of data and introduce extra complexity. For this reason, BIKG uses a Gene Target concept which is denoted by the gene ID, but merges the representation of the whole gene-transcript-protein subgraph.

To maintain the consistency of the model, concepts of the BIKG ULO ontology are mapped to the standard domain ontologies (such as Uberon [31] or Gene Ontology (GO) [1]) and, indirectly, to the foundational Basic Formal Ontology (BFO) ^4^. Moreover, the BIKG ULO defines ontological constraints on classes and properties to support quality assurance, most importantly, property domain/range and class disjointness: after building the integrated graph, automated tests verify whether the data conforms to the restrictions.

### 3.6 Identity resolution

One of the most typical and frequent data quality issue faced when ingesting heterogeneous data is *node duplication*, i.e. the same concept being represented in the graph by different node identifiers (e.g. HP:0030358 as Non-small cell lung cancer and UMLS:C0007131 as Non-small cell lung carcinoma). Node duplication causes to data fragmentation that degrades the quality of the data as well as leads to user confusion and data distrust. Duplication occurs for several reasons: different data sets use different vocabularies; one of the main data sources is the literature data, where the input data is evolving (by changing vocabularies used to tag nodes, new publications). Some node types are less ambiguous than others, for example we only encounter several gene vocabularies (which still requires extensive work on alignment), but there are dozens of disease vocabularies found in the ingested data sets (aligning disease nodes is a common problem when building large biomedical knowledge graphs, as there are many popular coding systems, where conversion is often not trivial [44]).

Node duplication is resolved by computing a stable set of mappings and merging the identified duplicate nodes. Each *duplicate node cluster* (set of nodes describing the same thing) is assigned a canonical, representative node, (e.g. for Gene nodes, Ensembl is preferred) and the node information of the other non-representative nodes (such as types, labels, other context fields) are merged into the canonical node. For example node type Side Effect is preserved, even if it is classified as a Disease in the final graph; this particular information can be used for graph completion where the existing edge data (e.g. *has_side_effect* edge type) does not already imply this fact.

Mappings are collected from public sources (e.g. data sets, research papers), licensed sources, as well as produced internally in collaboration with other teams. A stable set of mappings is produced as an iterative process: first all available mappings relating to the nodes of ingested sources are aggregated and the set of inferred mappings computed. Next the produced merges are manually examined (sampled) and tuned with the help of user feedback. Due to the challenges of manual curation, this focuses is on several high priority areas supporting the use cases. Mappings are retained for determining node merges during the graph build process (the node IDs in edges are updated using this table), but also for the purpose of adding data to the graph post build (as described in Section 3.1.3, users can add their own data to the produced graph using a a dedicated API).

### 3.7 Quality checking

Given the diversity of data sources and the volume of the data, data quality issues are inevitable. These are caused by either by imperfect data in the original sources (e.g. data entry errors, missing mappings etc) or by incorrect decisions taken by the data integration pipeline (e.g., a wrong canonical ID or label chosen during node merging). As noted by Stolios et al [44], several checks are required to be conducted in order to ensure the data quality of large biomedical knowledge graphs. This section first introduces the test framework (3.7.1) used in the pipeline, then it describes general data tests (3.7.2). Next it shows how node duplication is monitored (3.7.3), and finally it outlines how checking semantic constraints checking are implemented (3.7.4).

#### 3.7.1 Great Expectations

The knowledge graph build pipeline ingests and produces hundreds of different data files, such as parsed sources, intermediate build graph parts, different graph projections for distributing and accessing the graph (5.4), browsing (4.1), querying (via GraphDBs 4.2, Neo4j), and hosting graph analysis and content documentation. The *Great*

*Expectations* ^5^ *(GE)* data test framework is used in the graph build pipeline to validate the input and output data, configuration and other files. GE is a Python-based open-source library for validating, documenting, and profiling the data. GE generates a *data documentation website* (as illustrated by Figure 3) that helps to maintain data quality and improve communication about data between teams. GE supports several environments (Spark, Pandas, SQL) for working with columnar data, which makes it suitable for use on BIKG.

**Figure 3.**
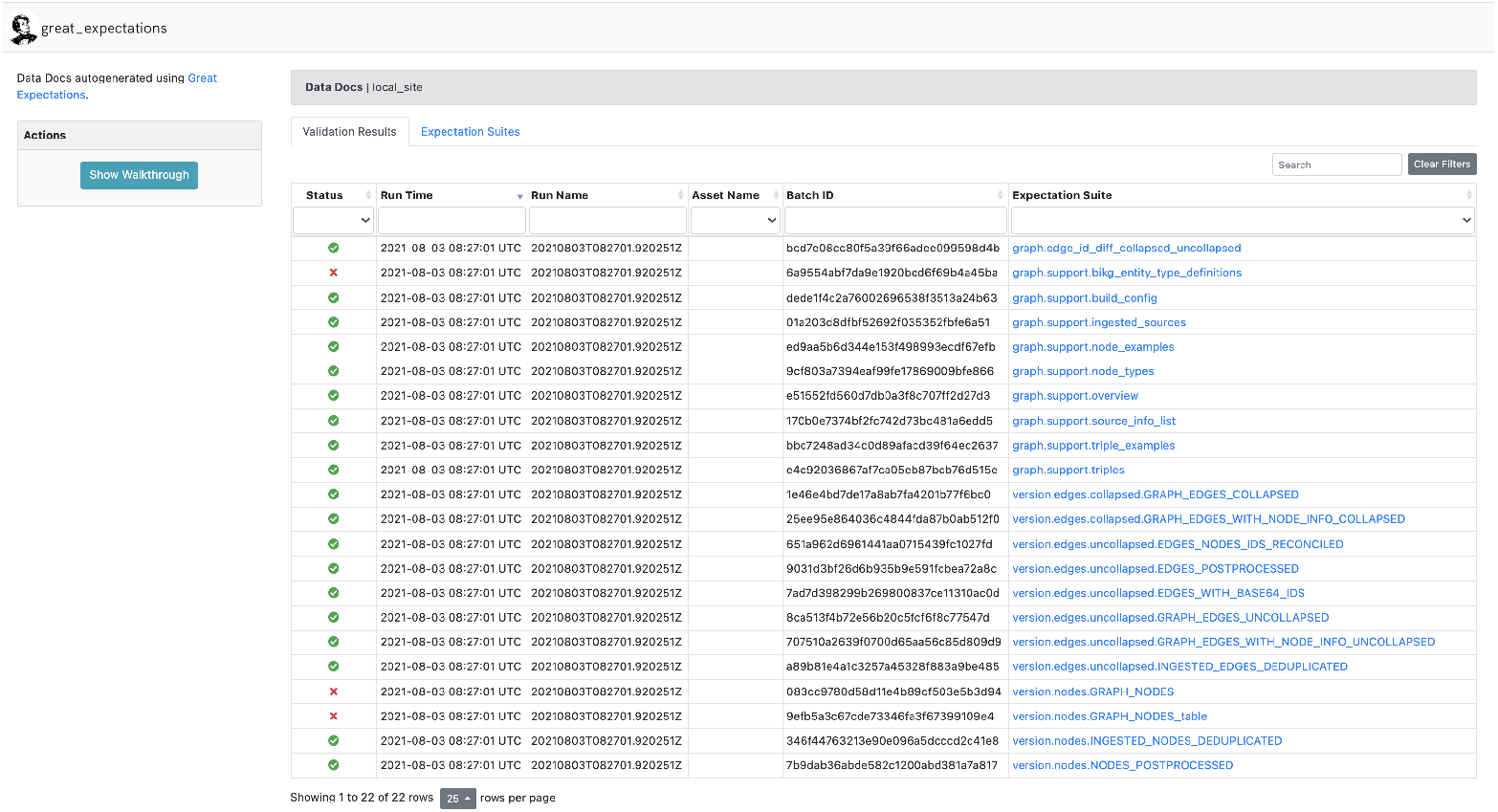
Great Expectations data documentation website

GE provides a number of predefined simple (and some more complex) data tests for validating on columnar data^6^. Tests such as *expect_column_values_to_be_in_set* can be used for checking the node *type* column content ULO conformance (see section 3.7.4). In addition, GE facilitates creating custom tests; for example ULO constraint checks (described in Section 3.7.4) require class subsumption reasoning, hence these are implemented as custom tests.

#### 3.7.2 Data tests

A variety of data tests are run on the graph to ensure its quality. Some tests are more general such as validating whether all node IDs referenced in the edge table are present in the node table, or more data specific such as node and triple existence, or validating the existence of certain nodes (along with their preferred label, specified typed, merged equivalent nodes). Data specific test cases typically focus on gene targets, diseases and compounds (see Section 3.5) in certain therapeutic areas; these test are typically hand curated.

#### 3.7.3 Node duplication check

In order to measure node duplication, node labels are compared for similarity. This can be challenging due to size (almost 11m nodes) hence heuristics are employed to reduce the search space (e.g. Gene Alterations are often wrongly named after the gene being altered using the gene symbol leading to duplication with the given gene node, therefore we can exclude these cases as false positives). In addition, duplication cases can are also reported by users using internal reporting channels and issue tracking software. Some of the node duplication analysis results, as well as user reports are used as proposed mappings and fed back into the graph, or used for expanding the data tests.

#### 3.7.4 ULO constraint check

The BIKO ULO (see Section 3.5) defines the class and property constraints used for validating the graph content. The ULO constraint checks ensure that: only permitted node and edge types are used, node types do not cause disjointness violations (see node “types” column), property domain and range violations are avoided and there are no restricted triple types. The checks are run on the tabular graph format, using Spark to materialise inferences. This is necessary due to scaling issues (memory requirements of triple stores), when using large RDF documents and running SHACL (Shapes Constraint Language, the W3C standard language for describing and validating RDF graphs [25]). Moreover, the RDF format is only one of many projections (different materialisation) of the graph, where the *main* projection is a set of parquet tables.

### 3.8 Pipeline overview

BIKG is built with a reproducible data pipeline that runs in the cloud and is capable of scaling to deal with very large amount of data. Figure 4 shows the main steps of the graph build pipeline.

**Figure 4.**
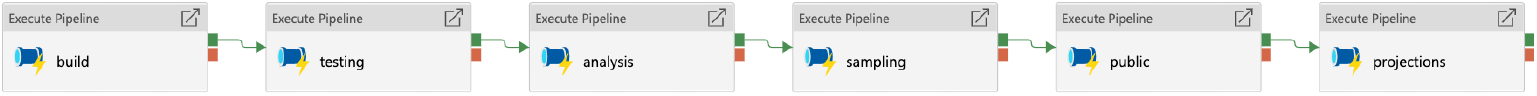
Graph build pipeline main steps

**Figure 5.**
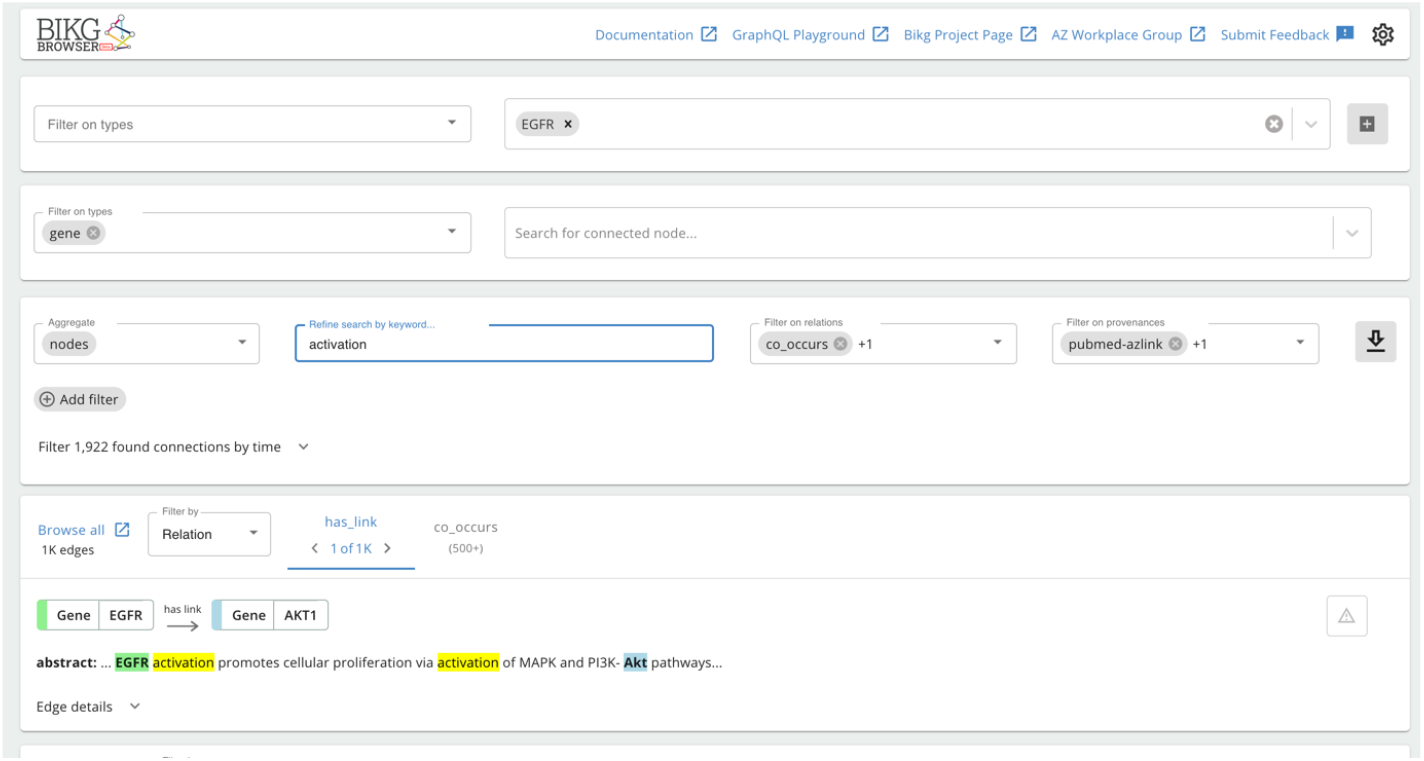
The BIKG Browser user interface, showing summary of the edges related to the EGFR gene.

- **Build**: the set of sources specified in the configuration are loaded and merged into one table according to data type: node, mapping or edge. Each table has a set of standardised columns (3.2) and potentially other columns that are merged into a single table containing all columns (this results in a sparse table as the different node types have different contextual data). Node deduplication (see Section 3.6), and edge compression (see Section 3.4) takes place in this step.
- **Testing**: syntactic and semantic tests are run on the graph, as described in Section 3.7.
- **Analysis**: the graph content is analysed and documented. This includes computing simple graph metrics (node, edge and triple type breakdowns, counts), and producing small but representative examples of nodes and edges, for visual inspection.
- **Sampling**: due to the large size of the full graph, several small samples are produced to facilitate testing and bench marking.
- **Public**: this step produces a subset of the full graph. As a future work, we intend to open source a large portion of BIKG, that is composed of publicly available data.
- **Projections**: this step creates several projections of the graph. The different projections contain all, or most of the graph data but materialised in different file formats to serve different purpose (e.g. RDF for loading in a triple store, CSV for loading in Neo4j etc.).

Parsing of raw data sources takes place outside of the graph build pipeline. The raw data comes in many different formats (CSV, TSV, parquet, RDF, multiline JSON and JSON-LD etc.) and obtained from internal and external data base API calls and data dumps. The graph is rebuilt each time a new dataset is added, removed or updated; therefore it is important to manage graph size and subsequently the required graph build process time and compute effort.

### 3.9 Tech stack overview

The graph build takes place the cloud. All graph data is stored in columnar format. In order to be able handle the large number, potentially billions of, edges (rows) and hundreds of features (columns) the pipeline uses *Spark* [51]. Most of the data is stored as parquet files to reduce size and improve the processing with Spark. Files are stored securely on an *Blob storage* [6] that complies with AstraZeneca data handling requirements for various types of data (public, licensed, company confidential and company restricted levels).

The code base is mostly implemented in Scala in order to optimise execution time. Python is also used in the project to allow for using a wider range of libraries, and to not limit developers and users to Scala. *Azure Datafactory* [24] is used as the ETL pipeline orchestrator. The compute clusters are managed by *Databricks* [12], where Python, Scala or R code is executed. In addition, the Databricks notebook environment is used for prototyping and collaborative development. *Azure Pipelines* [10] are used for running code base CI/CD, graph version and source analysis and data testing, creating and distributing releases and documentation.

## 4 Data access and exploration

Depending on the user profile and the task, different ways of interacting with the graph are required: e.g., keyword search, structured queries, or feeding data to machine learning models. There are different data storage, management, and access solutions that are optimised to tackle these interaction modalities: for example, indices for keyword search or graph databases for structured queries. However, it is difficult to select a single best solution that would be equally suitable for all of them. For this reason, after the BIKG graph is built, it is made available to the end users via several different access routes.

### 4.1 Browser

The BIKG browser is a web interface tool giving the users the capabilities to search and browse the graph data in a convenient form. At the back-end side it is powered by a custom Elasticsearch^7^ index which captures BIKG nodes as a set of multi-field documents, enabling faceted fuzzy keyword search. The browser index provides a GraphQL^8^ API enabling expressive queries for nodes and edges. This GraphQL API both serves the web UI as well as gives the end users the ability to retrieve data directly from their scripts. The web UI is optimised for two tasks: retrieving the nodes and edges using a combination of keyword queries and logical filters and browsing the graph data by following incoming and outgoing edges. While this interaction mode provides a convenient entry point for graph exploration, it does not allow specifying expressive structural queries by defining conditions over a whole subgraph pattern. This requirement is satisfied by using graph databases.

### 4.2 Graph DBs

In order to enable complex structural queries over the BIKG graph data, the graph is converted into a format suitable for loading into a graph database. The graph databases market includes two main streams of development: RDF triple stores and property graphs. Originally, RDF triple stores were primarily optimised for processing complex structural SPARQL queries, while property graphs commonly focused on the graph traversal algorithms. For this reason, BIKG graph gets exported into two formats suitable for these two different modalities: RDF^9^ and Neo4j^10^ (CSV).

### 4.3 Columnar storage dumps

Given that the graph data is mainly used for data analytics and ML tasks, it is important to make the data easily accessible from Python scripts. To this end, providing a data dump in a compact format together with the corresponding Python API making the data consumption straightforward was found to be both faster and more convenient than retrieving the data by querying a graph database. For this reason, the graph is also released using the partitioned Apache Parquet format^11^. This format is then consumed by the Python API described in Section 5.4.

## 5 Graph Machine Learning on BIKG

### 5.1 Knowledge Graph Embeddings

Recent years have seen an increase in methods that learns to encode the structural information of graphs as feature vectors. In this approach, called *representational learning*, each node in the graph is mapped to a point in a low-dimensional vector space, such that the graph topology is preserved. The node feature vectors, or knowledge graph embeddings, can then be used as input to downstream machine learning tasks.

A part of the BIKG project has been to provide embeddings, both for use in internal ML models and as a resource for researchers across AZ. Using Azure ML, we developed an automated pipeline for creating and analysing (see Section 5.2) embeddings after a new graph version is released.

For a large-scale graph like the BIKG, a computationally efficient way of calculating graph embeddings is not only desirable but necessary. To this end, we have used DGLKE [52], a tool for generating embeddings developed by Amazon Web Services. DGLKE enables distributed computing of embeddings across multiple GPUs.

### 5.2 Benchmarks

With each new graph release, we run a set of benchmark tasks to gain insight into how changes to the graph anatomy affect the performance of downstream machine learning models. The benchmarks include common biomedical machine learning tasks such as predicting Gene-Disease association and Protein-Protein Interaction (PPI). For each benchmark, the graph is reduced in size to only contain relevant entities, e.g., Gene and Disease nodes for Gene-Disease link prediction. Then, using 5-fold cross-validation, node embeddings are trained with RESCAL [33] using 80% of the edges and evaluated on the remaining 20%.

For evaluation, the node embedding vectors are concatenated for each pair of nodes that share an edge in the training set. These pairs are labelled as true examples. In addition, an equal amount of concatenated embeddings are created for nodes that do not share an edge. These pairs are labelled as false examples. Finally, a downstream machine learning model, XGBoost [7], is trained to distinguish between true and false examples. The performance of the XGBoost model tells us how well the embeddings capture the proximities in the graph.

To explore how the embeddings perform on different tasks with different data sources and how performance varies across graph releases, we created a Streamlit ^12^ application (Figure 6) displaying benchmark results from the ten latest graph releases.

**Figure 6.**
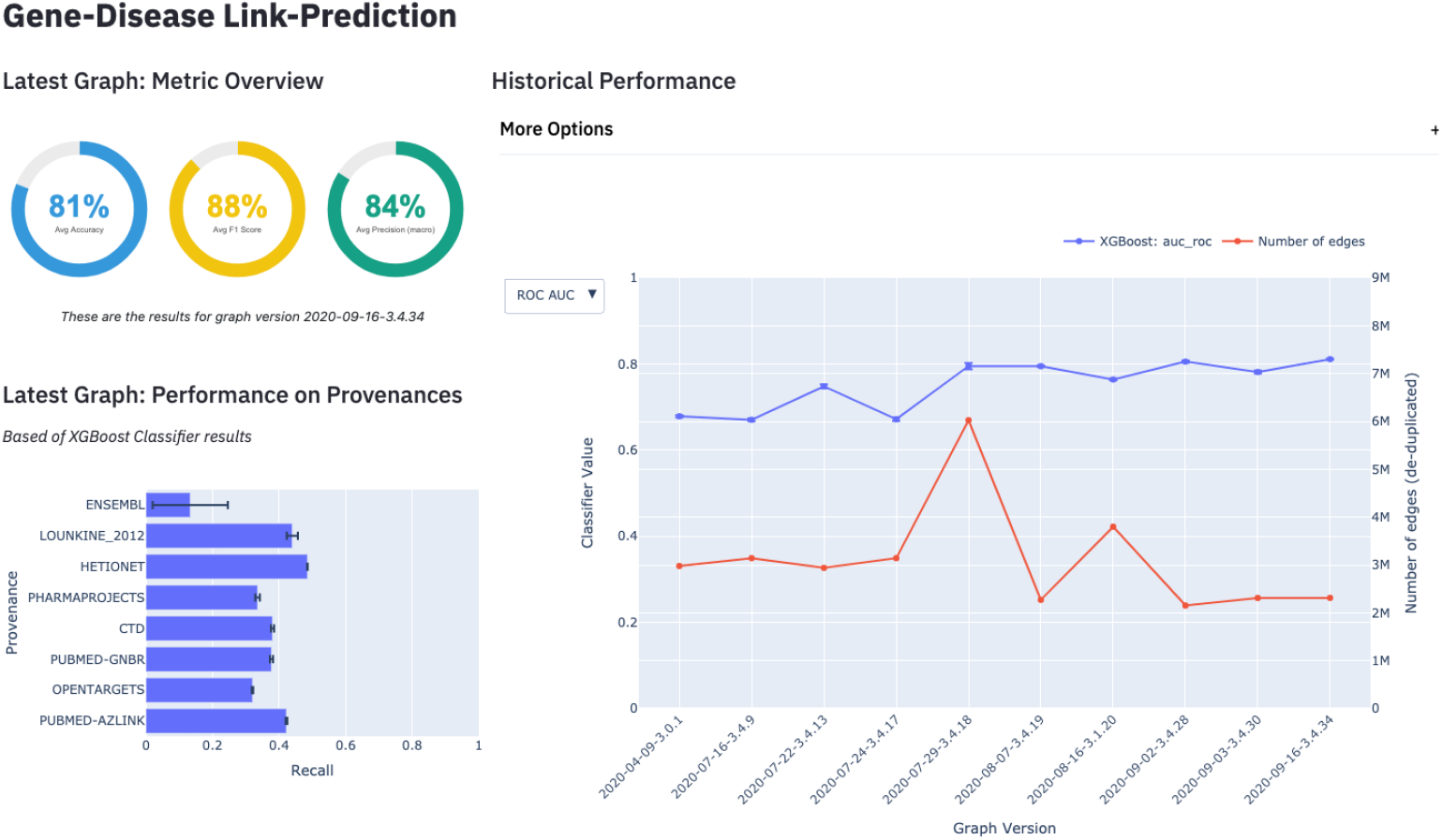
BIKG Benchmark Streamlit Application

**Figure 7.**
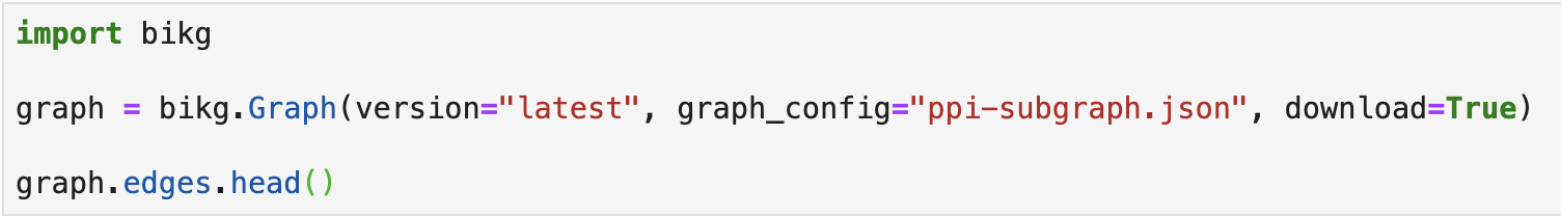
BIKG Python library usage

**Figure 8.**
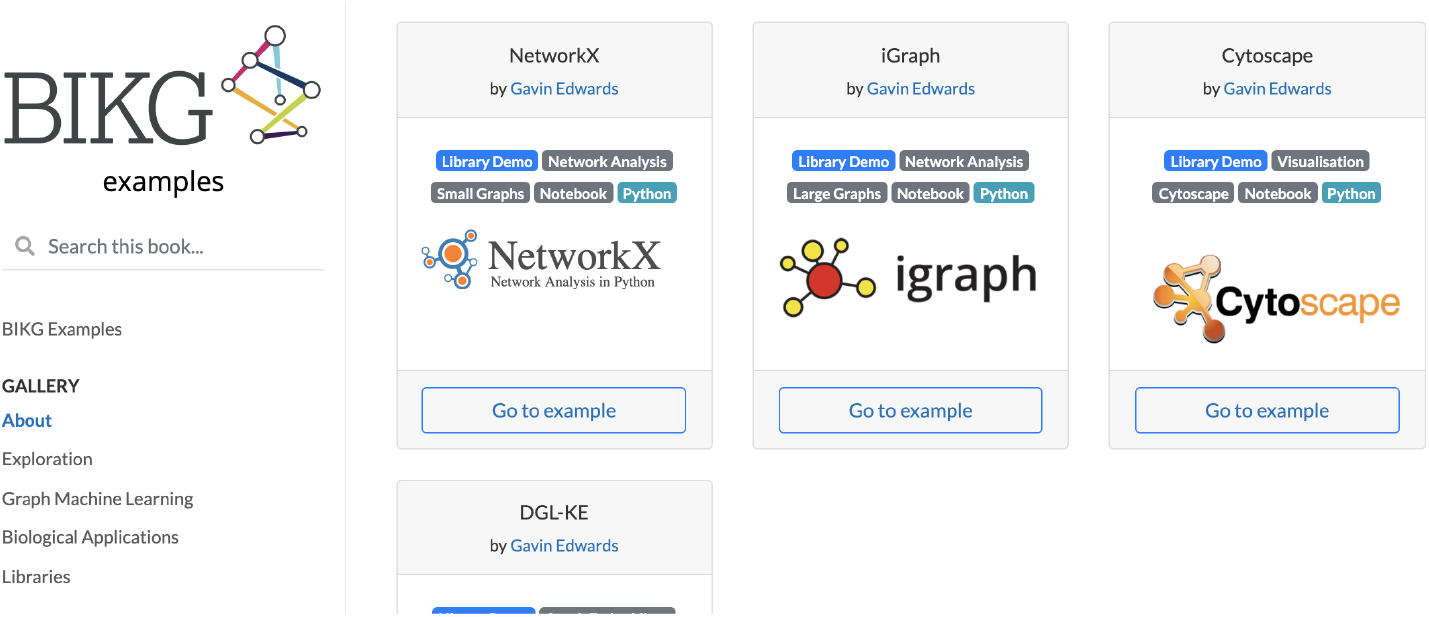
BIKG example tutorials

### 5.3 Graph Features

Graph-derived features, such as degree, betweenness, or clusterisation contain rich information about entities and edges in a graph. This information is valuable for downstream tasks, such as features for Machine Learning.

Generating graph derived features on the scale of BIKG is complex. Traditional tools, such as NetworkX [17], iGraph [8] and Spark GraphFrames [9] fail to scale. After experimenting with tools it was found that GPU powered network analysis was far superior. Therefore GPU backed cuGraph library [45] was used to generated graph derived features. It runs most algorithms on a single GPU, for more demanding algorithms such as betweenness, it leverages multi-GPU batch processing. By using this approach graph features can be computed in a few hours on the full BIKG.

Graph features are provided to users via two methods:

- By distributing pre-made features on the full BIKG graph.
- By providing the users with the example graph features notebook to create graph features on their own subgraphs.

The end result of this is it improves downstream tasks performance and enables users to quickly leverage the rich graphical information within the BIKG structure for their use cases.

### 5.4 Python Library

To maximise the impact of BIKG in Data Science and Machine Learning, it’s important to make accessing the data intuitive and quick. This enables users to experiment with the graph with minimal effort.

We developed a Python package that is focused on loading the subset of BIKG data a user wants to train on. We chose Python due to the wider ecosystem and its integration to other tools. The scope of the library is to make loading and cleaning subgraphs and their extra data as quick as possible. Once loaded, the data is stored in Pandas DataFrames, giving users instant familiarity and maximum portability to many other libraries. The package is accompanied by a documentation website with quick starts, tutorials, information and API explanations.

The same python library allows users to add their own data and enrich BIKG. This feature is especially important for sensitive data, since the data never has to leave the users’ environment.

Distributing a large dataset to end users via a python library can be challenging. We were able to maintain acceptable speed and efficiency by partitioning the data, leveraging fast network connectivity and caching datasets near the users environments. Another challenge was limiting the scope of the library: users often requested additional pre-built features. We chose to actively limit the scope of our Python package to load and filter the data, rather than trying to address all downstream tasks with a single software library. We felt the integrative approach would have ultimately limited the users, given the rapidly evolving landscape of external downstream libraries. Instead, we produced a large set of example Jupyter notebooks ^13^ which demonstrate common graph tasks and that users could customise freely.

The end result is users can quickly access, explore and apply BIKG data without any help.

### 5.5 Tutorial Notebooks

Quickstart example notebooks are made available, that show users how to load BIKG via the python library then perform other tasks. There are a variety of examples. Some demonstrate loading BIKG into other libraries, such as NetworkX [17], Cytoscape [43] or cuGraph [45]. Others demonstrate how to apply BIKG, for example to do link prediction or drug re-positioning.

The aim is to teach users how to approach graphs and do tasks. Instead of limiting users by wrapping around libraries, we provide them examples they can customise and extend without restriction.

Currently we include tutorials for

- exploratory data analysis with python and R
- extracting a disease subgraph
- examine literature trends
- filter edge types based on metadata, year or Mesh clinical term
- finding the best subgraph
- link prediction
- node classification
- explainable learning on graphs

We also include template workflows for the most common tasks we face when supporting decisions in the drug discovery pipeline

- unsupervised recommendations with graph features (triaging hits)
- protein function prediction
- drug repositioning
- target identification
- side effect prediction for combinations
- gene ranking using tensor factorisation
- target prediction with time slicing

There are high barriers getting started with BIKG due to unfamiliarity with (knowledge) graphs, the size of BIKG, new libraries and how to approach graph problems. The examples reduce the barrier of entry by bootstrapping new users and sharing the teams knowledge.

## 6 Use cases

The goal of the BIKG is to serve as an asset for domain specialists in providing non-trivial insights for solving use case problems. In this section we just briefly reference two BIKG use cases: target identification and re-ranking of CRISPR screen hits.

### 6.1 Target identification

When searching for novel gene targets, interpreting vast quantities of data across multiple experiments is a time-consuming, but necessary procedure. The challenge is to reduce time and effort and provide a more unbiased approach to identify novel targets. To address this issue, we built a decentralised pipeline for integrating the experimental data important to the user. Once ingested, these data, along with information derived from BIKG, produces a ranked list of potential gene targets for the chosen disease area.

We employed the pipeline to rank potential targets for Chronic Obstructive Pulmonary Disease, Asthma and Lung Cancer among others with significant internal success. We also tested the pipeline on public data by trying to predict novel targets before their publication. For this, we used the data published before 2015 as a training set and used the newly discovered relevant targets for the disease of interest published between 2015 and 2020 as a test set. The model could predict 16 out of 95 relevant new targets appearing in the following 5 years. We observe higher performance when we use more data types and focus on disease-specific subgraphs.

### 6.2 Re-ranking of CRISPR screen hits

Acquired drug resistance is a major factor making development of lasting cancer treatments difficult. One strategy for determining key drivers of acquired resistance is based on functional genomic screens, such as CRISPR screens [28].

The output of a CRISPR screen identifies thousands of potential targets. To narrow down the list to the most promising genes the scientists go through a lengthy and laborious procedure of manual validation, often prone to individual bias. There is a critical need for a standardised approach that integrates diverse types of knowledge to quickly identify valuable hits for experimental validation. We accelerated this decision-making process with a recommendation system built on top of BIKG.

The recommendation engine uses a multi-objective Pareto optimisation producing a Pareto front of potentially optimal solutions not dominated by others.

Resistance-specific hybrid feature set includes NLP-derived and graph-derived clinical and preclinical features. Assigning different importance to features, the user can re-rank the results according to the specific use case requirements. The results of this work are described in detail in [15].

## 7 Conclusion

The focus of the BIKG project is to build an internal knowledge graph leveraging both internal and external BIKG data, such that it can be used for applying machine learning algorithms and gaining new insights. This approach has demonstrated its value in several use cases, such as CRISPR recommendation and target identification.

Integrating various public datasets served as a backbone on top of which internal use case-specific data was added and contextualised, giving enough information for the trained algorithms to function.

Beyond the positive impact from achieving these goals, our experience has shown that such a knowledge graph also provides secondary benefits at the organisation level:

- Reduced time for data preparation work: common data preparation tasks like aligning the data formats and identity resolution are taken care when building the graph, which leaves scientists more time for actual research.
- Improved quality: dedicated APIs and templates enable quick addition of user data on demand as well as quick feedback from stakeholders.
- Reduced costs: the core graph as well as common task solutions are reusable across various use cases as well as across organisation units.

In our future work we are going to concentrate on the adaptation of the BIKG graph to new use cases and improving its suitability for novel machine learning techniques, such as graph neural networks, reinforcement learning, and explainable AI.

## Acknowledgments

We’d like to thank

## Footnotes

1 https://pubmed.ncbi.nlm.nih.gov/

2 https://parquet.apache.org/

3 https://en.wikipedia.org/wiki/N-Triples

4 http://basic-formal-ontology.org/

5 https://docs.greatexpectations.io/docs/

6 https://docs.greatexpectations.io/docs/reference/glossary_of_expectations

7 https://www.elastic.co/elasticsearch/

8 https://graphql.org/

9 https://www.w3.org/RDF/

10 https://neo4j.com/

11 https://parquet.apache.org/

12 https://streamlit.io/

13 https://jupyter.org/documentation

